# Modality-specific and multisensory mechanisms of spatial attention and expectation

**DOI:** 10.1101/847921

**Authors:** Arianna Zuanazzi, Uta Noppeney

## Abstract

In our natural environment, the brain needs to combine signals from multiple sensory modalities into a coherent percept. While spatial attention guides perceptual decisions by prioritizing processing of signals that are task-relevant, spatial expectations encode the probability of signals over space. Previous studies have shown that behavioral effects of spatial attention generalize across sensory modalities. However, because they manipulated spatial attention as signal probability over space, these studies could not dissociate attention and expectation or assess their interaction.

In two experiments, we orthogonally manipulated spatial attention (i.e., task-relevance) and expectation (i.e., signal probability) selectively in one sensory modality (i.e., primary modality) (experiment 1: audition, experiment 2: vision) and assessed their effects on primary and secondary sensory modalities in which attention and expectation were held constant.

Our results show behavioral effects of spatial attention that are comparable for audition and vision as primary modalities; yet, signal probabilities were learnt more slowly in audition, so that spatial expectations were formed later in audition than vision. Critically, when these differences in learning between audition and vision were accounted for, both spatial attention and expectation affected responses more strongly in the primary modality in which they were manipulated, and generalized to the secondary modality only in an attenuated fashion. Collectively, our results suggest that both spatial attention and expectation rely on modality-specific and multisensory mechanisms.

## Introduction

Spatial attention is a top-down mechanism that is critical for the selection of task-relevant information. It facilitates perception (e.g., faster reaction times, greater accuracy) of signals presented at the attended location (Carrasco, 2011; van Ede et al., 2012). By contrast, spatial expectation (signal probability) facilitates perception by encoding the statistical structure of the environment (Summerfield & Egner, 2009; Rohenkohl et al., 2014). In everyday life, spatial attention and expectation are closely intertwined. For instance, observers often allocate attentional resources to locations in space where events are more likely to occur (Summerfield & Egner, 2009; Feldman & Friston, 2010 for further discussion within the predictive coding framework).

Importantly, in our natural multisensory environment our brain is constantly exposed to auditory and visual signals. This raises the critical question of whether allocation of attention and encoding of signal probability are performed in a modality-specific fashion or interactively across sensory modalities. Previous research has suggested that spatial attention relies on cognitive resources that are partially shared across sensory modalities (Eimer & Driver, 2001; Wahn & König 2015, 2017). For instance, Spence & Driver (1996) manipulated spatial attention by presenting signals with a higher probability in the attended relative to unattended hemifield in one modality only (i.e., primary modality). They showed behavioral facilitation for signals presented at the attended location not only for the primary modality (e.g., audition) but also for the secondary modality (e.g., vision) in which spatial attention was not explicitly manipulated. Likewise, neuroimaging studies showed increased activations for signals presented at the attended location not only in the primary modality in which attention was manipulated but also in the secondary modality (Eimer & Schröger, 1998; Eimer, 1999; Macaluso et al., 2002; Santangelo et al., 2009; Zuanazzi & Noppeney, 2019). Crucially, in this past work the attentional effects were greater in the primary than in the secondary modality (Spence & Driver, 1996; Mondor & Amirault, 1998). The attenuated generalization across sensory modalities suggests that attentional resources are not supramodal, but partially shared (Driver & Spence, 1998). However, this past research conflated attention and expectation by manipulating spatial attention via probabilistic spatial cues or changes in signal probability (Posner, 1980; Spence & Driver, 1996, 1997; Macaluso et al., 2002; Kincade et al., 2005; Bressler et al., 2008; Santangelo et al., 2009). Notably, the Posner probabilistic cuing paradigm shifts observers’ attention via spatial cues that indicate whether a target is, for instance, more likely to appear in the left or right hemifield. Likewise, manipulating not only categorically whether the cue is valid or invalid but also its validity (e.g., 100% vs 60% valid) (Vossel et al., 2006; Doricchi et al., 2010; Macaluso & Doricchi, 2013) does not enable the dissociation of spatial attention and expectation. Observers should allocate their attentional resources more to their left hemifield when presented with a cue that indicates with a probability of 1 rather than 0.6 whether the target is likely to be presented in the left hemifield.

Thus, the first question of this study is whether spatial attention and/or expectation generalize across sensory modalities to a similar extent, when they are manipulated independently. In the most extreme case, they may be modality-specific (i.e., no generalization) or amodal (i.e., complete generalization). A recent neuroimaging study, for instance, suggested that spatial attention relies mainly on frontoparietal cortices for both primary and secondary modalities, while spatial expectations are formed in sensory systems selectively for the primary modality (Zuanazzi & Noppeney, 2019). As a consequence, we would expect spatial signal probability to be encoded selectively for the primary modality and to generalize to a secondary modality only to a limited degree.

A second unresolved question is whether the generalization across sensory modalities depends on whether attention and expectation are manipulated in the auditory or visual modalities as primary manipulation modality - i.e., on the direction of cross-sensory generalization. Previous studies of multisensory attention have indeed shown asymmetric multisensory generalization, depending on which modality was manipulated as primary modality (Ward at al., 2000; Greene et al., 2001; Molholm et al., 2007). Moreover, one may expect differences in cross-sensory generalization from vision to audition and vice versa because spatial representations and expectations are encoded differently in audition and vision. Visual and auditory systems encode space via different reference frames (i.e., eye-centered vs head-centered) and representational formats. In the visual system, spatial location is directly encoded in the sensory epithelium and later in a place code, i.e., via retinotopic organization of primary and higher order visual cortices (e.g., Sereno et al., 1995; Maier & Groh, 2009). In the auditory system, spatial locations are computed from binaural and monoaural cues in the brain stem and are represented in a hemifield code in primary auditory cortices (e.g., Lauter et al., 1985; Maier & Groh, 2009). Further, in everyday life under normal lighting conditions, vision usually provides more reliable spatial information than audition and therefore often dominates spatial perception (Spence & Driver, 1997; Aller et al., 2015; Odegaard et al., 2015; Rohe & Noppeney, 2015a, 2015b. 2016, 2018; Aller & Noppeney, 2019; Jones et al., 2019; Meijer et al., 2019). As a result, we would expect the generalization of spatial attention and expectation to depend on whether attention and expectation are manipulated primarily in vision or audition.

Third, in everyday life spatial expectations are formed when observers implicitly learn the statistical structure of their multisensory environment such as the probability of signals occurring at a particular location. Because spatial information is encoded less reliably in audition than vision, this learning may be faster in vision than in audition. Thereby spatial expectations may also be affected by multisensory processes in perceptual learning (e.g., Kim et al., 2008; Batson et al., 2011). For instance, previous studies suggested that perceptual learning in temporal discrimination tasks generalizes across sensory modalities (Warm et al., 1975; Nagarajan et al., 1998; Meegan et al. 2000; Bratzke et al., 2012; Bueti et al., 2012, 2014; but see Lapid et al., 2009). This study will investigate how observers dynamically form spatial expectations by learning signal probability over time (Crist et al., 1997).

In two experiments (within participants), we orthogonally manipulated spatial attention and expectation selectively in one sensory modality as primary modality (experiment 1: audition, experiment 2: vision). Crucially, to dissociate spatial attention and expectation we did not use a probabilistic cuing paradigm. Instead, we manipulated observers’ spatial attention in the primary (experiment 1: audition, experiment 2: vision) modality by instructing them to attend and respond to, e.g., auditory targets selectively in their left but not right hemifield. In addition, we manipulated the relative frequency of auditory stimuli in the left (e.g., 30%) and right (e.g., 70%) hemifield. Because observers need to respond only to targets in their left hemifield, they should ideally allocate all attentional resources to this task-relevant hemifield irrespective of stimulus frequency. Moreover, observers had to respond to all stimuli in their secondary (e.g., visual) modality irrespective of the hemifield in which they were presented. These visual stimuli were presented equally often in both hemifields. We then assessed the effects of spatial attention and expectation in the primary modality and how they generalize crossmodally to signals in the secondary sensory modality, in which spatial attention and expectation were not explicitly manipulated (experiment 1: vision, experiment 2: audition).

If attention, expectation and decision making rely on modality-specific processing streams (i.e., without any multisensory interplay), we would expect as null-hypothesis that the attention and expectation manipulations in the primary modality would not affect response times to stimuli in the secondary modality. Hence, the alternative hypothesis is that both spatial attention and expectations rely on mechanisms that are partially shared across sensory modalities and the generalization of spatial attention and expectation depends on whether they are manipulated in audition or vision as primary modalities (Spence & Driver, 1997; Ward at al., 2000; Greene et al., 2001; Molholm et al., 2007; Aller et al., 2015; Odegaard et al., 2015; Rohe & Noppeney, 2015a, 2015b. 2016, 2018; Aller & Noppeney, 2019; Jones et al., 2019; Meijer et al., 2019).

## Materials and Methods

### Participants

Twenty-eight healthy subjects (19 females; mean age = 25.57 years; 24 right-handed) participated in the study (experiment 1 and experiment 2, within participants). The sample size was determined based on previous studies that investigated attention/expectation (Doherty et al., 2005; van Ede et al., 2012; Beck et al., 2014; Rohenkohl et al., 2014) and/or multisensory integration (Spence & Driver, 1996, 1997; Eimer et al., 2004; Santangelo et al., 2008; Krumbholz et al., 2009; Mengotti et al., 2018; Zuanazzi & Noppeney, 2018).

All participants had normal or corrected to normal vision and reported normal hearing. All participants provided written informed consent and were naïve to the aim of the study. The study was approved by the local ethics committee of the University of Birmingham (Science, Technology, Mathematics and Engineering (STEM) Ethical Review Committee) and the experiment was conducted in accordance with these guidelines and regulations.

### Stimuli and Apparatus

Spatial auditory stimuli of 100 ms duration were created by convolving bursts of white noise (with 5 ms onset and offset ramps) with spatially selective head-related transfer functions (HRTFs) based on the KEMAR dummy head of the MIT Media Lab (http://sound.media.mit.edu/resources/KEMAR.html). Visual stimuli (‘flashes’) were white discs (radius: 0.88° visual angle, luminance: 196 cd/m2) of 100 ms duration presented on a grey background. Both auditory and visual stimuli were presented at ±10° of horizontal visual angle along the azimuth (0° of vertical visual angle). Throughout the entire experiment, a fixation cross was presented in the center of the screen.

Prior to the beginning of the study, participants were tested for their ability to discriminate left and right auditory stimuli on a brief series of 20 trials. They indicated their spatial discrimination response (i.e., ‘left’ vs ‘right’) via a two-choice key press (group mean accuracy was 99% ± 0.4% [across subjects mean ± SEM]).

During the experiment, participants rested their chin on a chinrest with the height held constant across all the participants. Auditory stimuli were presented at approximately 72 dB SPL, via HD 280 PRO headphones (Sennheiser, Germany). Visual stimuli were displayed on a gamma-corrected LCD monitor (2560 x 1600 pixels resolution, 60 Hz refresh rate, 30” Dell UltraSharp U3014, USA), at a viewing distance of approximately 50 cm from the participant’s eyes. Stimuli were presented using Psychtoolbox version 3 (Kleiner et al., 2007; www.psychtoolbox.org), running under Matlab R2014a (Mathworks Inc., Natick, MA, USA) on a Windows machine. Participants’ responses were recorded via one key of a small keypad (Targus, USA). Throughout the study, participants’ eye-movements and fixation were monitored using Tobii Eyex eyetracking system (Tobii EyeX, Tobii, Sweden, ~60 Hz sampling rate).

### Study overview: rationale and analysis strategy

This study included two experiments. Each experiment conforms to a four-factorial design. Because the two experiments were performed within the same participants, the study as a whole could also be treated as a five factorial within-subject experiment. However, because (1) experiment 1 and 2 were completed on different days, (2) experiment 1 was a replication of our previous study and (3) the understanding of results of five factorial designs is rather complex, we will initially analyse each of the two experiments separately.

The separate analyses of experiments 1 and 2 allow us to address our first question, i.e., whether spatial attention and expectations rely on modality-specific or at least partially shared mechanisms. While experiment 1 is intended to replicate our findings reported in our previous research (Zuanazzi & Noppeney, 2018), experiment 2 is intended to extend them and demonstrate that this pattern of results does not depend on whether audition or vision is used as a primary manipulation modality. Moreover, showing the same profile across the two experiments also resolves the ambiguity of our previous research, in which a smaller spatial expectation effect for the secondary (i.e., visual) modality in experiment 1 could potentially be explained by differences in sensory modality rather than attenuated cross-sensory generalization.

In a second step we will directly combine data from experiment 1 and 2 to address our second question, i.e., whether the spatial expectation effect generalizes differently from audition to vision than from vision to audition. To address this question, we need to compare the expectation effects for auditory and visual stimuli in the attended hemifield between the two experiments (i.e., this question cannot be addressed by any of the two experiments alone).

Please also note that the Design and Procedure section mostly overlaps with that of our previous paper (Zuanazzi & Noppeney, 2018) to enable the reader to quickly compare our different studies and obtain a convergent picture across all results.

### Design and Procedure

In two experiments, participants were presented with auditory and visual stimuli in their left and right hemifields. To manipulate spatial attention, they were instructed to respond to stimuli in the primary sensory (e.g., auditory) modality selectively in one (i.e., task-relevant) hemifield and ignore stimuli in the task-irrelevant hemifield. Moreover, we manipulated observers’ spatial expectations by presenting stimuli in the primary sensory modality with different probabilities in the task-relevant and irrelevant hemifields. In their secondary (e.g., visual) modality, observers had to respond to all stimuli that were presented equally often in both hemifields (Fig. 1A and 1B). Experiment 1 investigated the effect of auditory spatial attention and expectation on detection of auditory (i.e., primary modality) and visual (i.e., secondary modality) targets using a 2 (auditory spatial attention: left vs right hemifield) x 2 (auditory spatial expectation: left vs right hemifield) x 2 (stimulus modality: auditory vs visual) x 2 (stimulus location: left vs right hemifield) factorial design. Hence, experiment 1 manipulated spatial attention and expectation selectively in audition and assessed their direct effects on auditory stimulus processing and indirect generalization to visual stimuli. In experiment 2 primary and secondary modality were reversed (i.e., primary modality: vision; secondary modality: audition); design and procedural details were otherwise comparable to experiment 1. For the data analysis we pooled over stimulus locations (left/right) leading to a 2 (auditory spatial attention: attended vs unattended) x 2 (auditory spatial expectation: expected vs unexpected) x 2 (stimulus modality: auditory vs visual) factorial design.

**Figure 1:**
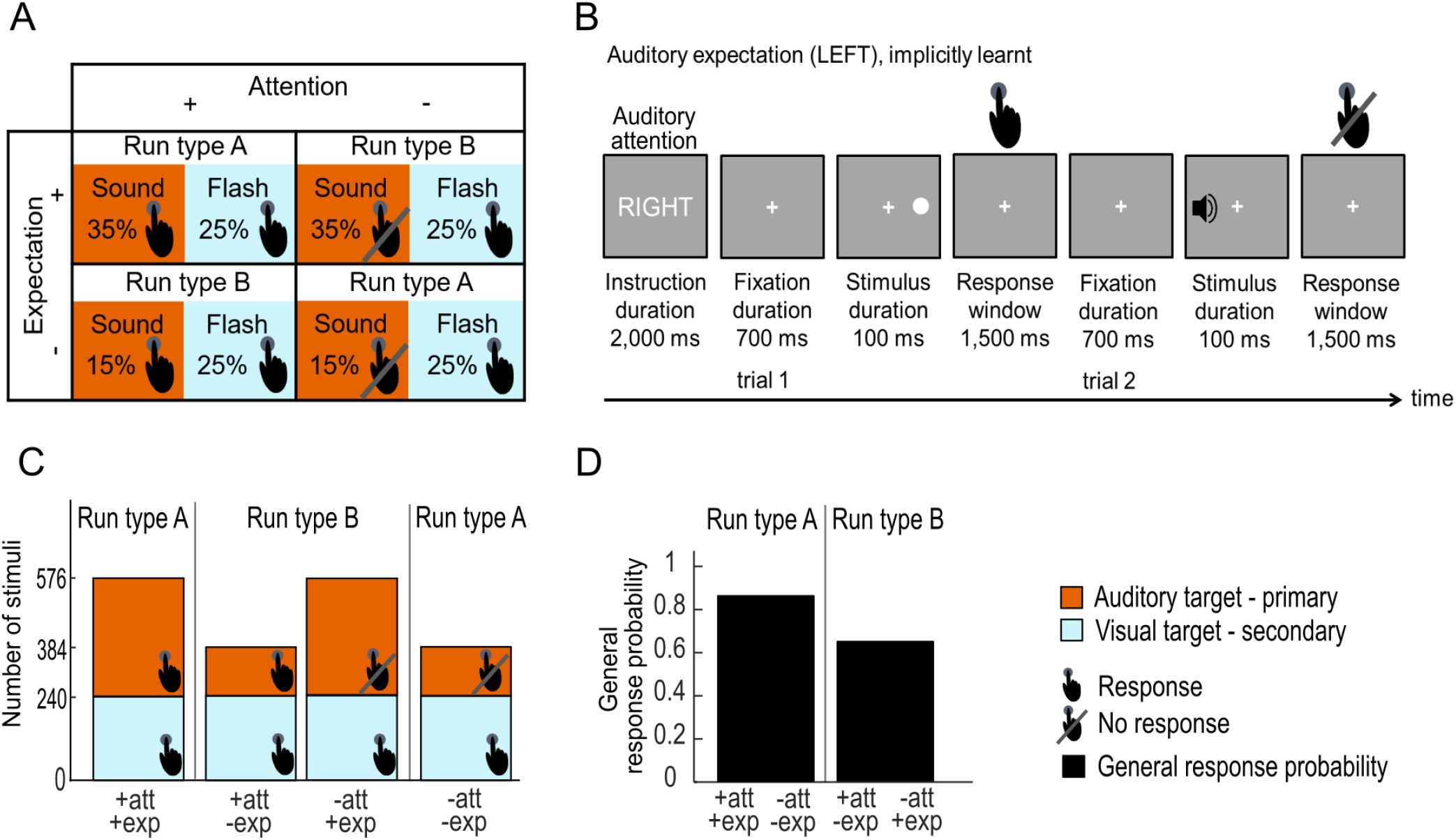
Design and example trials of experiment 1 (audition to vision) **A.** Experiment 1: auditory spatial attention and expectation (i.e., signal probability) were manipulated in a 2 (auditory modality - dark orange, vs visual modality - light blue) x 2 (attended hemifield vs unattended hemifield) x 2 (expected hemifield vs unexpected hemifield) factorial design. For illustration purposes, stimulus locations (left/right) were collapsed. Presence vs absence of response requirement is indicated by the hand symbol, spatial signal probability manipulation is indicated by the %. **B.** Experiment 1: example of two trials in a session where auditory stimuli were presented with a probability of 0.7 in the left hemifield and 0.3 in the right hemifield. At the beginning of each run (i.e., 80 trials), a cue informed participants whether to attend and respond to auditory signals selectively in their left or right hemifield throughout the entire run. On each trial participants were presented with an auditory or visual stimulus (100 ms duration) either in their left or right hemifield. They were instructed to respond to auditory stimuli only in the attended hemifield and to all visual stimuli irrespective of the hemifield as fast and accurately as possible with their index finger. The response window was limited to 1500 ms. Participants were not explicitly informed that auditory signals were more likely to appear in one of the two hemifields. Instead, spatial expectation was implicitly learnt within a session (i.e., day). **C.** Experiment 1: number of auditory (dark orange) and visual (light blue) trials in the 2 (attended vs unattended hemifield) x 2 (expected vs unexpected hemifield) design (pooling over left/right stimulus location). Presence vs absence of response requirement is indicated by the hand symbol. The fraction of the area indicated by the ‘Response’ hand symbol pooled over the two bars of one particular run type (e.g., run type A) represents the response related expectation (i.e., general response probability: the overall probability that a response is required on a particular trial); general response probability is greater for run type A (85%), where attention and expectation are congruent, than for run type B (65%), where attention and expectation are incongruent, as indicated in **D**. Note. Design and procedure of experiment 2 were comparable to that of experiment 1, with the only difference that vision was the primary modality and audition was the secondary modality. In other words, in experiment 2 attention and expectation were manipulated selectively in vision.

Spatial attention was manipulated for the primary modality as task-relevance, i.e., the requirement to respond to an auditory (experiment 1) or a visual (experiment 2) target in the left vs right hemifield. Prior to each run a cue (duration: 2000 ms) informed the observer whether to respond to targets in either their left or right hemifield.

Spatial expectation was manipulated as spatial signal probability for signals in the primary modality across experimental sessions that were performed on different days. Auditory (i.e., primary modality in experiment 1) or visual (i.e., primary modality in experiment 2) signals were presented with a ratio of 2.33/1 (i.e., 70%/30%) in the expected/unexpected hemifield. Observers were not informed about those probabilities but learnt them implicitly. Importantly, spatial attention and expectation were not directly manipulated in the secondary modality, allowing us to assess their cross-sensory generalization. As a result, participants needed to respond to all visual targets that were presented with equal probabilities in their spatial hemifields in experiment 1 (i.e., ratio 1/1 in the expected/unexpected hemifields) (Fig. 1A and 1C). Likewise, they had to respond to all auditory targets that were presented with equal probabilities in experiment 2.

Each experiment included two sessions (i.e., spatial expectation left vs right on different days). Hence, subjects participated in the two experiments on four days separated by at least 2 to a maximum of 10 days: 2 sessions for experiment 1 and 2 sessions for experiment 2 = 4 sessions in total for each participant. Each session included 12 attention runs. Runs were of two types: in run type A (Fig. 1A, 1C and 1D) spatial attention and expectation were congruent (i.e., spatial attention was directed to the hemifield with higher stimulus frequency); in run type B spatial attention and expectation were incongruent (i.e., spatial attention was directed to the hemifield with less frequent stimuli). The overall probability to respond (i.e., response probability) was greater when attention and expectation were congruent and directed to the same hemifield (85%, runs of type A) than when they were directed to different hemifields (65%, runs of type B) (Fig. 1D).

The order of experiments 1 vs 2 and of expectation sessions (i.e., left vs right) was counterbalanced across participants; the order of attention runs (i.e., left vs right) was counterbalanced within and across participants and the order of stimulus locations (i.e., left vs right) and stimulus modalities (sound vs flash) was pseudo-randomized within each participant. Brief breaks were included after every two attention runs to provide feedback to participants about their performance accuracy (averaged across all conditions) in the target detection task and about their eye-movements (i.e., fixation maintenance).

Overall, each experiment included 80 trials x 12 attention runs (6 runs of type A and 6 runs of type B, duration: 3 mins/run) x 2 expectation sessions = 1920 trials in total (and 3840 for the whole study). Specifically, each run type included i. 336 targets presented in the expected hemifield (pooled over left and right) and 144 targets in the unexpected hemifield (pooled over left and right) for the primary modality and ii. 240 targets presented in the expected hemifield and 240 targets in the unexpected hemifield (pooled over left and right) for the secondary modality. For further details see Fig. 1C which shows the absolute number of trials for each condition and run type.

Each trial (SOA: 2300 ms) included three time windows: i. fixation cross alone (700 ms duration), ii. brief sound or flash (stimulus duration: 100 ms) and iii. fixation cross alone (1500 ms as response interval, see Fig. 1B). Participants responded to the targets in the primary modality presented in the attended hemifield and to all targets in the secondary modality irrespective of hemifield via key press, with their index finger (i.e., the same response for all auditory and visual stimuli) as fast and accurately as possible (Fig. 1B).

Prior to each session, participants were familiarized with the stimuli in brief practice runs (with equal spatial signal probability) and trained on target detection performance and fixation (i.e., a warning signal was shown when the disparity between the central fixation cross and the eye-data samples exceeded 2.5 degrees).

After the final session (i.e., experiment 1 for 14 participants and experiment 2 for the other 14 participants), participants indicated in a questionnaire whether they thought the stimuli in the primary modality were presented more frequently in one of the two spatial hemifields. Ten out of 14 participants in experiment 1 and 11 out of 14 participants in experiment 2 correctly identified the expectation manipulation in the primary modality. Moreover, 13 out of 14 participants in experiment 1 and 13 out of 14 participants in experiment 2 correctly reported that stimuli in the secondary modality were presented with equal probabilities across the two hemifields. These data suggest that the majority of participants were aware of the manipulation of signal probability at least at the end of the fourth session.

## Data analysis

### Eye movement: exclusion criteria

We excluded trials where participants did not successfully fixate the central cross based on a dispersion criterion (i.e., distance of fixation from subject’s center of fixation, as defined in calibration trials, > 1.3 degrees for three subsequent samples; Blignaut, 2009). Our eyetracking data confirmed that participants successfully maintained fixation in both experiments with only a small number of trials to be excluded (experiment 1: excluded auditory response trials 1.8% ± 0.5% [across subjects mean ± SEM]; excluded visual response trials 1.7% ± 0.5% [across subjects mean ± SEM]; experiment 2: excluded visual response trials 2.7% ± 1% [across subjects mean ± SEM]; excluded auditory response trials 2.7% ± 0.9% [across subjects mean ± SEM]).

### Response time analysis - separately for experiments 1 and 2

We initially analysed response times separately for primary and secondary modalities and independently for experiments 1 and 2.

The response time (RT) analysis was limited to trials with RT within the 1500 ms response window and was performed after pooling over stimulus location (left/right).

For the primary modality (i.e., experiment 1 = audition, experiment 2 = vision), subject-specific median RT were entered into a two-sided paired-sample t-test with spatial expectation (expected vs unexpected stimulus) as factor (observers did not respond to targets in the primary modality in the ‘unattended’ hemifield). Moreover, subject-specific False Alarm rates (FA) for the ‘unattended’ hemifield were entered into a non-parametric two-sided Wilcoxon Signed Rank tests with spatial expectation (expected vs unexpected stimulus) as factor. We used non-parametric tests, because False Alarms rates are bounded between 0 and 1 and therefore not normally distributed. For the secondary modality (i.e., experiment 1 = vision, experiment 2 = audition), subject-specific median response time were entered into a 2 (spatial attention: attended vs unattended stimulus) x 2 (spatial expectation: expected vs unexpected stimulus) repeated measures analysis of variance (ANOVA).

For both experiments 1 and 2, the mean hit rates were very high (> 99% in all conditions, Table 1), indicating that participants accurately performed the detection task. Because of the absence of a substantial number of misses, hit rates were not further analyzed.

**Table 1:** Group mean hit rates, false alarm (FA) rate and reaction times (RT) and for each stimulus modality in each condition where a response was given for experiment 1 (primary modality: audition, secondary modality: vision) and experiment 2 (primary modality: vision, secondary modality: audition). In experiment 1, participants responded only to attended auditory targets (and to all visual targets); in experiment 2 participants responded only to attended visual targets (and to all auditory targets). Standard errors (SEM) are given in parentheses.

|  | **Auditory modality** |  |  |  | **Visual modality** |  |  |  |
| --- | --- | --- | --- | --- | --- | --- | --- | --- |
| **Experiment 1** | +att<br>+exp | +att<br>-exp | -att<br>+exp | -att<br>-exp | +att<br>+exp | +att<br>-exp | -att<br>+exp | -att<br>-exp |
| Hit rate (%)<br>(SEM) | 99.5<br>(0.19) | 99.5<br>(0.15) | / | / | 99.7<br>(0.07) | 99.7<br>(0.07) | 99.3<br>(0.16) | 99.4<br>(0.13) |
| FA rate (%)<br>(SEM) | / | / | 4.4<br>(0.8) | 7.1<br>(0.1) | / | / | / | / |
| RT (ms)<br>(SEM) | 599.7<br>(20.1) | 616.2<br>(19.2) | / | / | 514.4<br>(16.9) | 522.1<br>(16.9) | 583.8<br>(18.5) | 574.4<br>(18.9) |
| **Experiment 2** | +att<br>+exp | +att<br>-exp | -att<br>+exp | -att<br>-exp | +att<br>+exp | +att<br>-exp | -att<br>+exp | -att<br>-exp |
| Hit rate (%)<br>(SEM) | 99.8<br>(0.08) | 99.9<br>(0.03) | 99.2<br>(0.22) | 99.4<br>(0.18) | 99.7<br>(0.09) | 99.7<br>(0.09) | / | / |
| FA rate (%)<br>(SEM) | / | / | / | / | / | / | 3.2<br>(0.5) | 8<br>(1) |
| RT (ms)<br>(SEM) | 527.4<br>(20.2) | 542.1<br>(20) | 605.2<br>(25.7) | 581.3<br>(24.2) | 526.3<br>(17.9) | 561.8<br>(19.9) | / | / |

### Response time analysis - combined for experiments 1 and 2

To compare effects of primary vs secondary modality, we compared the response times in the attended hemifield (averaged across expected and unexpected hemifields) for auditory and visual stimuli across the two experiments in a 2 (stimulus modality: audition vs vision) x 2 (manipulation: primary/direct vs secondary/indirect modality) repeated measures ANOVA.

Next, we investigated whether the effect of spatial expectation (i.e., expected vs unexpected) in the attended hemifield (no response was required for stimuli in the primary modalities presented in the unattended hemifield) depended on i. whether targets were presented in the primary or secondary modalities (i.e., the extent to which spatial expectations generalize across the senses) and ii. the multisensory generalization direction, from audition to vision and from vision to audition (i.e., whether spatial expectations generalize differently depending on whether audition or vision is the primary modality). Hence, we first computed the difference in median RT (ΔRT_Exp_) between unexpected and expected stimuli presented in the attended hemifield (which corresponds to ΔRT_Exp_ between attended stimuli in run type B and run type A) for targets in each experiment, yielding four conditions: i. auditory targets as primary modality (experiment 1), ii. visual targets as secondary modality (experiment 1), iii. visual targets as primary modality (experiment 2), iv. auditory targets as secondary modality (experiment 2). ΔRT_Exp_ were entered into a 2 (multisensory generalization direction: audition to vision vs vision to audition) x 2 (manipulation: primary/direct vs secondary/indirect modality) repeated measures ANOVA. Please note that the interaction then reflects the difference between targets in the auditory and visual modality.

### Time course of response times – combined for experiment 1 and 2

Finally, we assessed how these effects of multisensory generalization direction and direct/indirect manipulation evolved over time. For this, we computed the difference in median RT (ΔRT_Exp_) between unexpected and expected stimuli, as in the previous analysis, but now separately for the first and second half of the experiment (i.e., one half = 430 trials). Each half contained the data from 6 subsequent attention runs (3 runs of type A and 3 runs of type B) for each expectation condition. ΔRT_Exp_ (or each half) were entered in a 2 (multisensory generalization direction: audition to vision vs vision to audition) x 2 (manipulation: primary/direct vs secondary/indirect modality) x 2 (time: first vs second half of the experiment) repeated measures ANOVA. Figure 2D shows the across subjects’ mean (±SEM) RT separately for each of the 2 attention runs (1 of type A and 1 of type B) (orange and blue circles) as a more fine-grained temporal characterization of the effects of expectation over time. An additional analysis using this more fine-grained temporal division replicated the results reported in this manuscript where we separated the data into halves.

**Figure 2:**
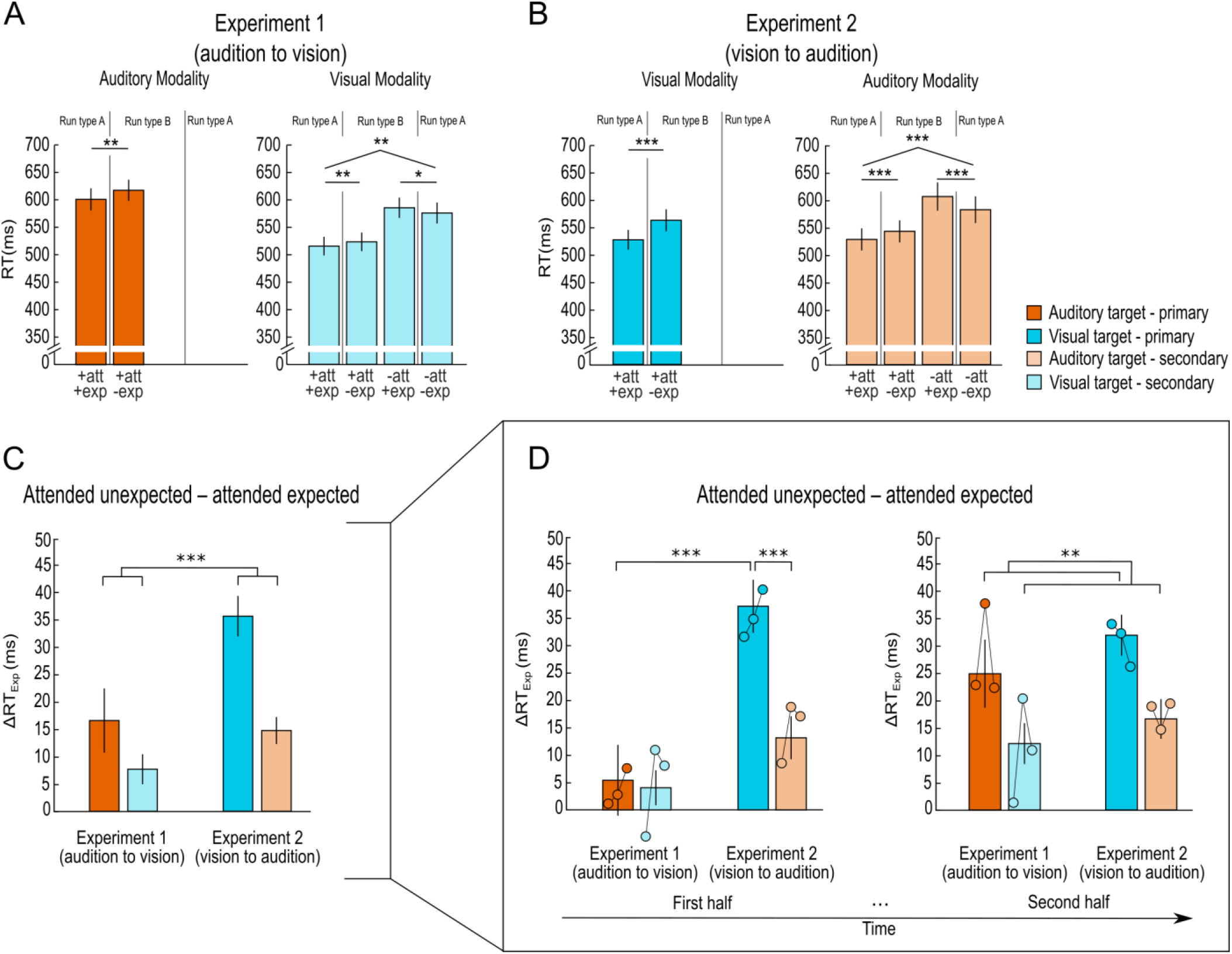
Behavioural results of experiment 1 and 2. Bar plots represent across subjects’ mean (±SEM) RT for each of the six conditions with response requirements for experiment 1 (primary modality: audition; secondary modality: vision, **A**) and 2 (primary modality: vision; secondary modality: audition, **B**), pooling over left/right stimulus location. Overall slower RT are observed for runs type B than runs type A, which reflects differences in general response probability (see Fig. 1D) **C.** Bar plots represent across subjects’ mean (±SEM) ΔRT for effects of spatial expectation (attended unexpected - attended expected hemifield) in the primary (dark bars) and secondary modalities (light bars) for experiment 1 and 2. **D.** Effects of response probability (attended unexpected - attended expected hemifield) over time (i.e., first and second half: bars; consecutive sets of 2 attention runs: circles) for audition and vision as primary (dark bars) or secondary (light bars) modality. Brackets and stars indicate significance of main effects and interactions. \**p* < 0.05; ** *p* < 0.01; *** *p* < 0.001. Audition: orange; vision: blue.

For all analyses we assessed the assumptions of normality using the Shapiro–Wilk test (Shapiro and Wilk, 1965). When normality was violated, we evaluated the main effects of attention, expectation and their interactions in the factorial design using permutation testing with 2^28^ permutations (Nichols & Holmes, 2002). Because in these cases permutation tests replicated the results of the initial ANOVAs, we only report the results of the ANOVAs for consistency.

## Results

### Generalization of attention and expectation effects across modalities - separately for experiment 1 and 2

In experiment 1, participants responded to auditory targets presented in their attended hemifield and to all visual targets. In experiment 2, participants responded to visual targets presented in their attended hemifield and to all auditory targets.

We observed qualitatively similar effects across experiments 1 and 2. Table 1 shows RT (across participants’ mean ± SEM) for targets in the auditory and visual modalities for the two experiments.

For the primary modality, the two-sided paired-sample t-tests showed significantly faster RT in the attended hemifield, when this hemifield was expected than unexpected (experiment 1, auditory modality: *t*(27) = −2.83, *p* = 0.009, Cohen’s *d_av_* [95% CI] = −0.16 [−0.27, −0.04]; experiment 2, visual modality: *t*(27) = −9.62,*p* < 0.001, Cohen’s *d_av_* [95% CI] = −0.35 [−0.47, −0.23]) (Table 1, Fig. 2A and 2B). Moreover, the Wilcoxon Signed Rank tests showed significantly greater FA in the unattended hemifield, when stimuli in this hemifield were unexpected than expected (experiment 1, auditory modality: W = 52, *p* = 0.001, r [95% CI] = −0.72 [−0.87, −0.45]; experiment 2, visual modality: W = 2, *p* < 0.001, r [95% CI] = −0.99 [−1, −0.98]) (Table 1).

For the secondary modality, the 2 (attended vs unattended) x 2 (expected vs unexpected) repeated measures ANOVAs revealed a significant main effect of attention (experiment 1, visual modality: *F*(1, 27) = 72.08, *p* < 0.001, *η_p_*^2^ [90% CI] = 0.73 [0.55, 0.80]; experiment 2, auditory modality: *F*(1, 27) = 36.91, *p* < 0.001, *η_p_*^2^ [90% CI] = 0.58 [0.34, 0.70]). Results showed that participants responded faster to targets presented in their attended than unattended hemifields.

Moreover, a significant crossover interaction between attention and expectation was observed (experiment 1, visual modality: *F*(1, 27) = 10.09, *p* = 0.004, *η_p_*^2^ [90% CI] = 0.27 [0.06, 0.46]; experiment 2, auditory modality: *F*(1, 27) = 44.10, *p* < 0.001, *η_p_*^2^ [90% CI] = 0.62 [0.40, 0.73]) (Table 1, Fig. 2A and 2B). The simple main effects showed that participants responded significantly faster to targets in their attended hemifield, when this hemifield was expected than unexpected (experiment 1, visual modality: *t*(27) = −2.81, *p* = 0.009, Cohen’s *d_av_* [95% CI] = −0.08 [−0.15, −0.02]; experiment 2, auditory modality: *t*(27) = −5.96, *p* < 0.001, Cohen’s *d_av_* [95% CI] = −0.14 [−0.19, −0.08]). This significant simple main effect demonstrates that the effects of spatial attention and expectation generalized from primary to secondary modalities, where neither attention nor expectation were explicitly manipulated. By contrast, participants responded significantly more slowly to targets in the secondary modality in the unattended hemifield, when this hemifield was expected than unexpected (experiment 1, visual modality: *t*(27) = 2.56, *p* = 0.016, Cohen’s *d_av_* [95% CI] = 0.09 [0.02, 0.17]; experiment 2, auditory modality: *t*(27) = 5.53,*p* < 0.001, Cohen’s *d_av_* [95% CI] = 0.18 [0.10, 0.26]) (Table 1, Fig. 2A and 2B). We suggest that simple main effects for expectation show opposite directions because of response inhibition. In the attended hemifield observers need to respond to the stimuli in the primary modality. Hence, if stimuli from the primary modality are frequent (i.e., expected) in the attended hemifield, observers need to respond on a large percentage of trials. By contrast, in the unattended hemifield observers should not respond to the stimuli in the primary modality. Hence, if stimuli in the primary modality are frequent (i.e., expected) in the unattended hemifield, observers need to inhibit their response on a large percentage of trials. This explanation is also supported by the increase in FA for the primary modality in the unattended hemifield when stimuli in this hemifield are unexpected relative to expected (see above and Fig. 1D). Collectively, the response times and FA rates suggest that, in runs in which observers need to respond to many stimuli, because the stimulus frequency is high in the task-relevant/attended hemifield, observers will make more false alarms to stimuli of the primary modality and respond faster to stimuli of the secondary modality in the unattended hemifield. We can explain this profile in decision making models in which observers need to accumulate evidence to a threshold. An increase in the percentage of trials that require a response may then be reflected either in a shift of the starting point closer to the decisional boundary or in a lower decisional boundary (Gold & Shadlen, 2001).

### The effects of primary vs secondary modality – combined for experiment 1 and 2

To assess the effect of primary vs secondary modality unconfounded by differences between auditory vs visual modality, we directly compared the response times in the attended hemifield (averaged across expected and unexpected hemifields) for auditory and visual stimuli across the two experiments. The 2 (stimulus modality: audition vs vision) x 2 (manipulation: primary/direct vs secondary/indirect modality) repeated measures ANOVA revealed a significant main effect of stimulus modality (*F*(1, 27) = 40.14, *p* < 0.001, *η_p_*^2^ [90% CI] = 0.60 [0.37, 0.72]), showing faster RT for visual than auditory targets; of manipulation (*F*(1, 27) = 151.80, *p* < 0.001, *η_p_*^2^ [90% CI] = 0.85 [0.74, 0.89]), showing overall faster RT for secondary than primary modality. Moreover, we observed a significant interaction between stimulus modality and manipulation (*F*(1, 27) = 6.79, *p* = 0.015, *η_p_*^2^ [90% CI] = 0.20 [0.02, 0.39]), showing that observers were significantly faster responding to stimuli in the secondary than primary sensory modality predominantly in experiment 1 (primary = auditory, secondary = visual, *t*(27) = 11.67,*p* < 0.001, Cohen’s *d_av_* [95% CI] = 0.93 [0.63, 1.22]).

Collectively, these results suggest that observers responded faster to stimuli in their secondary modality than in their primary modality. As expected, this effect was more sensitively revealed for auditory stimuli that were associated with slower response times. These effects can be explained by the fact that observers needed to respond to all stimuli in the secondary modality irrespective of hemifield. By contrast, they first needed to discriminate whether signals were presented in the left or right hemifield when responding to stimuli in the primary sensory modality. Because the spatial reliability is lower for auditory than visual signals in our study, these effects were more prominent for auditory stimuli.

### The effects of spatial expectation - combined for experiment 1 and 2

In the previous analysis we showed that the effects of expectation on response times generalized from the primary to the secondary modality. Next, we directly compared the effects of spatial expectation across the two experiments, in which either audition was the primary and vision the secondary modality or vice versa. The 2 (multisensory generalization direction: audition to vision vs vision to audition) x 2 (manipulation: primary/direct vs secondary/indirect modality) repeated measures ANOVA revealed a significant main effect of manipulation (*F*(1, 27) = 18.03, *p* < .001, *η_p_*^2^ [90% CI] =0.40 [0.16, 0.56]), showing overall greater expectation effects for primary than secondary modalities (dark vs light bars in Fig. 2C). Moreover, we observed a significant main effect of crossmodal generalization direction (*F*(1, 27) = 20.71, *p* < .001, *η_p_*^2^ [90% CI] = 0 .43, [0.19, 0.59]), with a greater expectation effect (i.e., greater ΔRT_Exp_) for vision to audition than for audition to vision (experiment 2 vs experiment 1 in Fig. 2C). Critically, this generalization effect may be greater from vision to audition than vice versa because observers learn signal probabilities and hence form spatial expectations faster when vision is the primary modality (dark blue bar in Fig. 2C). Alternatively, the expectation effect generalizes more effectively from vision to audition than vice versa (light orange bar in Fig. 2C).

### Time course of the effects of spatial expectation - combined for experiment 1 and 2

To disentangle between these two possibilities, we investigated how signal probability is learnt over time when audition (experiment 1) or vision (experiment 2) are the primary modality by repeating the previous analysis with the additional factor of time (i.e., first vs second half of experiment). The 2 (multisensory generalization direction: audition to vision vs vision to audition) x 2 (manipulation: primary/direct vs secondary/indirect modality) x 2 (time: first vs second half of the experiment) repeated measures ANOVA performed on ΔRT_Exp_ revealed significant main effects of multisensory generalization direction (*F*(1, 27) = 22.70, *p* < 0.001, *η_p_*^2^ [90% CI] = 0.46 [0.21, 0.61]), manipulation (*F*(1, 27) = 13.77, *p* < 0.001, *η_p_*^2^ [90% CI] = 0.34 [0.10, 0.51]) as well as a significant interaction between multisensory generalization direction x manipulation x time (*F*(1, 27) = 11.38, *p* = 0.002, *η_p_*^2^ [90% CI] = 0.29 [0.07, 0.48]).

We unpacked the 3-way ANOVA into two 2-ways ANOVAs for further analysis, one for each half of the experiment, with factors multisensory generalization direction (audition to vision vs vision to audition) and manipulation (primary/direct vs secondary/indirect modality).

For the first half of the experiment, our results revealed a significant main effect of multisensory generalization direction (*F*(1, 27) = 8.34, *p* = 0.008, *η_p_*^2^ [90% CI] = 0.24 [0.04, 0.42]), manipulation (*F*(1, 27) = 28.84, *p* < 0.001, *η_p_*^2^ [90% CI] = 0.52 [0.27, 0.65]) and a significant interaction between multisensory generalization direction and manipulation (*F*(1, 27) = 12.83, *p* = 0.001, *η_p_*^2^ [90% CI] = 0.32 [0.09, 0.50]) (left bar plot in Fig. 2D). Post-hoc comparisons indicated that ΔRT_Exp_ in the primary modality of experiment 2 (i.e., vision) were significantly greater than ΔRT_Exp_ in the primary modality of experiment 1 (i.e., audition) (*t*(27) = 5.72, *p* < 0.001, Cohen’s *d_av_* [95% CI] = 1.05, [0.59, 1.50], dark blue vs dark orange bars in the left bar plot of Fig. 2D), and greater than the effects of expectation in the secondary modality of experiment 2 (i.e., audition) (*t*(27) = 4.97, *p* < 0.001, Cohen’s *d_av_* [95% CI] = 1.03, [0.53, 1.51], dark blue vs light orange bars in the left bar plot of Fig. 2D). Moreover, the effects of expectation in the secondary modality of experiment 2 (i.e., audition) were significantly greater than those in the secondary modality of experiment 1 (i.e., vision) (*t*(27) = 2.14,*p* = 0.042, Cohen’s *d_av_* [95% CI] = 0.48, [0.02, 0.94], light orange vs light blue bars in the left bar plot of Fig. 2D).

For the second half of the experiment, our results only revealed a significant main effect of manipulation (*F*(1, 27) = 11.41, *p* = 0.002, *η_p_*^2^ [90% CI] = 0.30 [0.07, 0.48]) (right bar plot in Fig. 2D) but no significant main effect of multisensory generalization direction or significant interaction between multisensory generalization direction and manipulation was found.

To summarize, these results show that: (1) an effect of generalization direction (i.e., audition to vision vs vision to audition) was found only for the first half of the experiment. Here, the effects of expectation generalized crossmodally in an attenuated fashion only from vision to audition in experiment 2 (i.e., primary modality: vision) but no difference between audition and vision was found in experiment 1 (i.e., primary modality: audition). By contrast, in the second half of the experiment we did not observe an effect of multisensory generalization direction. Instead, the effects of expectation generalized from the primary to the secondary modality in an attenuated fashion similarly when vision or audition were the primary modality. As shown in figure 2D, this difference between first and second halves can be explained by the fact that observers form spatial expectations (i.e., learn signal probability over space) more slowly in audition than vision. Yet, once expectations are learnt in audition, the crossmodal generalization is comparable for audition and vision. In other words, the effect of generalization direction that we observed in our analysis that did not yet account for learning effects (i.e., our second analysis) can be explained away by the speed with which observers learn signal probabilities and form spatial expectations in their primary modality. In other words, signal probabilities are learnt faster in vision than audition, but once spatial expectations are formed, they generalize similarly from vision to audition and vice versa.

## Discussion

The current study investigated how observers allocate attention and form expectations (by learning signal probabilities) over space across audition and vision. We orthogonally manipulated spatial attention as response requirement and expectation as stimulus probability over space selectively in the primary modality and assessed their effects on behavioral responses to targets presented in the primary and secondary modalities. Across two experiments we alternated the assignment of vision and audition to primary or secondary modality. This allowed us to compare behavioral effects of spatial attention and expectation in audition and vision and their crossmodal generalization.

Regardless of sensory modality we observed a significant main effect of spatial attention for targets in the secondary modality, in which attention was not directly manipulated. Auditory spatial attention partially generalized to the visual modality and vice versa. These findings converge with a large body of behavioral and neuroimaging work suggesting that attentional resources are allocated interactively across the senses (Spence & Driver, 1996; Eimer & Schröger, 1998; Eimer, 1999; Macaluso et al., 2002; Santangelo et al., 2009; Zuanazzi & Noppeney, 2019).

Likewise, in the attended hemifield observers were faster at target detection in the primary and secondary modalities when the hemifield was expected than unexpected (i.e., high > low signal probability). Again, this response facilitation for expected (relative to unexpected) spatial locations were observed irrespective of whether vision or audition served as primary modality. Yet, in the unattended hemifield, in which we could assess effects of expectation only for the secondary modality, we observed the opposite pattern, i.e., observers were slower at target detection for expected than unexpected hemifields. Combining these two results, we observed a significant interaction between spatial attention and expectation, for both vision and audition as secondary modalities. In a previous study (Zuanazzi & Noppeney, 2018) we argued that this interaction profile between spatial attention and expectation is explained by attention and expectation jointly co-determining general response probability (i.e., the probability to respond regardless of the hemifield in which the signal is presented). More specifically, in runs in which attention and expectation are directed to the same hemifield (runs of type A, Fig. 1A, 1C and 1D), participants have to respond to 85% of trials of the entire run, but only to 65% trials in runs of type B in which attention and expectation are directed to different hemifields (i.e., general response probability, Fig. 1A, 1C and 1D). Hence, faster response times may result from an increase in alertness, arousal or motor preparation that is needed to respond on a large proportion of trials (i.e., attended/expected and unattended/unexpected conditions, run type A, Fig. 2A and 2B) (Mars et al., 2007; Bestmann et al., 2008). By contrast, in runs in which attention and expectation are directed to different hemifields (runs of type B, Fig. 1A, 1C and 1D), observers need to inhibit their response to the frequent stimuli of the primary modality in the expected hemifield and hence respond more slowly to targets in the secondary modality.

Critically, response probability does not depend on whether the signal is auditory or visual, but it is calculated as the probability that any signal is responded to. If the expectation effects result purely from amodal mechanisms (e.g., general alertness, arousal, motor preparation etc.) associated with changes in response probability, the expectation effects in the attended hemifield should be equal for primary and secondary modalities. By contrast, if expectations (i.e., auditory or visual signal probability) are formed at least partially in a modality-specific fashion, we should observe expectation effects (i.e., ΔRT_Exp_) that are greater for the primary modality (where expectation was explicitly manipulated) than for the secondary modality (i.e., an attenuated crossmodal generalization).

To arbitrate between these two hypotheses, we analyzed ΔRT_Exp_ in a 2 (multisensory generalization direction) x 2 (manipulation) repeated measures ANOVA. This analysis revealed a significant main effect of ‘manipulation’, i.e., whether the stimulus was presented in the primary modality (in which expectation was explicitly manipulated) or in the secondary modality. Consistent with additional modality-specific mechanisms of expectation, observers showed a greater expectation effect for targets in the primary than the secondary modality. These results strongly suggest that implicitly learned spatial expectations modulate perceptual decision making via both modality-specific and amodal response mechanisms.

This duality of modality-specific and amodal mechanisms converge with recent neuroimaging findings which showed effects of expectation selective for auditory stimuli as primary modality in auditory cortices and higher-order frontoparietal systems (Zuanazzi & Noppeney, 2019). Potentially, the activation increases in auditory cortices may reflect prediction error signals based on modality-specific expectations (Friston, 2005), while higher frontoparietal systems may be associated with additional response-related processes.

Surprisingly, the repeated measures ANOVA also revealed a main effect of multisensory generalization direction with greater effects in experiment 2 (i.e., vision to audition) than experiment 1 (i.e., audition to vision). Our time course analysis showed that this difference between experiments does not reflect genuine differences in the effectiveness with which spatial expectations are implicitly learnt and generalize from audition to vision and vice versa, but reflects differences in the speed with which spatial expectations are learnt in audition and vision. In the second half of the experiment in which the expectation effect in the primary modality is comparable between auditory (experiment 1) and visual (experiment 2) targets, we no longer observe a significant effect of multisensory generalization direction. In other words, observers are slower at learning spatial expectations (i.e., signal probability) in audition than in vision. But, once they have formed spatial expectations of comparable precision for audition and vision, they also generalize similarly from audition to vision and vice versa.

The difference in perceptual learning rates between vision and audition may result from how the brain forms spatial representations in vision and audition (Neumann et al., 1986). While visual cortices are retinotopically organized and hence directly represent spatial location in a place code (i.e., based on space, Sereno et al., 1995; Maier & Groh, 2009), primary auditory cortices are tonotopically organized (i.e., based on frequency, Lauter et al., 1985, Middlebrooks &, 1991; Maier & Groh, 2009). In audition, spatial locations are computed only indirectly from binaural amplitude and latency differences and from monoaural filtering cues. Moreover, visual objects tend to be more permanent across time, whereas source sounds are often transient and dynamic (Neisser, 1976; Neumann et al., 1986). Most importantly, in everyday life vision provides typically (i.e., under optimal lighting conditions) more reliable spatial information than audition (Dacey et al., 1992; Knudsen & Brainard, 1995; Stephen et al., 2002; Talsma et al., 2008; Mengotti et al., 2018; see also Molholm et al., 2007). In the current study, the high spatial reliability of the visual stimulus (i.e., a white disc) may also have contributed to the shorter time for participants to learn spatial signal probabilities and thus become aware of their manipulation in vision. Critically, participants’ awareness of such manipulation is evidenced by our questionnaires’ results. Alternatively, even in absence of explicit awareness, the distribution of events or targets across space could have been learnt faster for more reliable visual than auditory signals (Miller & Pachella, 1973; Jabar & Anderson, 2015). Conversely, in a paradigm that investigates temporal attention/expectation mechanisms, the high temporal resolution and precision of auditory signals (Shimojo & Shams, 2001) could facilitate learning of temporal probability in audition more than in vision and crossmodal effects could change accordingly. The existence of cross-modal effects of temporal attention is shown in previous studies investigating how attention is oriented to different points in time (e.g., Lange & Röder, 2006). However, while the effects of spatial and temporal attention were similar for auditory processing, they differed for visual and tactile modalities, suggesting the existence of modality specific mechanisms also in the temporal domain. To better understand the fine-grained temporal aspects of spatial expectation or signal probability learning across sensory modalities, future studies will need to characterize the time course of spatial learning across sensory systems.

So far, we have discussed that spatial expectations generalize only partially across the senses. One critical question is whether this partial generalization is generic or arises because the decisions rely on different processes in our paradigm. Most importantly, as we have indicated in the Results section, observers had to respond to stimuli in the primary modality only in one hemifield, but in the secondary modality in both hemifields. This experimental choice enabled us to assess the additive and interactive effects of spatial attention and expectation in both hemifields for the secondary modality. As a consequence, however, observers needed to determine the hemifield in which the stimulus occurred before making a response only for the primary modality. By contrast, they could respond non-discriminatively to all stimuli in the secondary modality. This difference in the decision-making process most likely explains that observers were faster to respond to sounds when audition was the secondary than the primary sensory modality. An outstanding question is whether this difference in the decision-making process can also explain the partial generalization of the expectation effects. In other words, would we observe more extensive or perhaps complete generalization across sensory modalities if both primary and secondary modalities rely on similar decision-making processes? Or even more fundamentally, can the differences in magnitude in the expectation effects for primary and secondary modality be explained by the fact that expectations influence spatial discrimination processes that are required only for responding to stimuli in the primary modality? To address this question, future studies may manipulate attention and expectation in both sensory modalities. For instance, they may present observers with auditory and visual stimuli in left and right hemifields. Auditory stimuli may occur mainly in the left and visual in the right hemifield. Importantly, observers will need to respond to auditory and visual stimuli only when they occur in one particular (e.g., left) hemifield, so that responses to both auditory and visual stimuli will require spatial discrimination between hemifields. However, as we have argued in a previous study, manipulating response requirement and spatial expectations orthogonally across sensory modalities may interact at the several levels by jointly specifying not only observers’ general response probability but also spatially selective response probabilities (Zuanazzi & Noppeney, 2018). Further, it is important to emphasize that these manipulations of spatial signal probability and response requirement over space operate bidirectionally from audition to vision and vice versa as well as from primary to secondary modality and vice versa, thus interpretational ambiguities may remain. Alternatively, complementary insights may be gained from neuroimaging research that can implicitly assess the multisensory generalization of neural representations linked with spatial expectations even when no response is required.

In summary, our results suggest that the brain allocates spatial attention and forms spatial expectation to some extent interactively across audition and vision (Eimer & Schröger, 1998; Eimer, 1999; Macaluso, 2010). With respect to spatial attention, our results corroborate previous research. With respect to spatial expectation, we show that they rely on modality-specific and amodal mechanisms. In support of modality-specific mechanisms we demonstrate that spatial expectations in the attended hemifield generalize from the primary to the secondary modality only in an attenuated fashion. In support of amodal response-related mechanisms, we demonstrate that, for both primary and secondary modalities, response times are closely related to the general response probability and associated processes of arousal, alertness and motor preparation. Critically, our learning analysis suggests that observers learn spatial probabilities more slowly in audition than vision, which may be related to their different spatial reliabilities. Once observers have formed comparable spatial expectations in audition, these generalize equally effectively from audition to vision as from vision to audition. In other words, our results demonstrate crossmodal interactions of perceptual learning (i.e., expectations building) in spatial perception but also show differences between sensory modalities in terms of the speed with which signal probabilities over space are learnt.

## Acknowledgments

We thank Catarina S. Ferreira and Remi Gau for their helpful comments on this manuscript.

